# Inferring Time-Varying Internal Models of Agents Through Dynamic Structure Learning

**DOI:** 10.1101/2024.09.30.615757

**Authors:** Ashwin James, Ingrid Bethus, Alexandre Muzy

## Abstract

Reinforcement learning (RL) models usually assume a stationary internal model structure of agents, which consists of fixed learning rules and environment representations. However, this assumption does not allow accounting for real problem solving by individuals who can exhibit irrational behaviors or hold inaccurate beliefs about their environment. In this work, we present a novel framework called Dynamic Structure Learning (DSL), which allows agents to adapt their learning rules and internal representations dynamically. This structural flexibility enables a deeper understanding of how individuals learn and adapt in real-world scenarios. The DSL framework reconstructs the most likely sequence of agent structures—sourced from a pool of learning rules and environment models—based on observed behaviors. The method provides insights into how an agent’s internal structure model evolves as it transitions between different structures throughout the learning process. We applied our framework to study rat behavior in a maze task. Our results demonstrate that rats progressively refine their mental map of the maze, evolving from a suboptimal representation associated with repetitive errors to an optimal one that guides efficient navigation. Concurrently, their learning rules transition from heuristic-based to more rational approaches. These findings underscore the importance of both credit assignment and representation learning in complex behaviors. By going beyond simple reward-based associations, our research offers valuable insights into the cognitive mechanisms underlying decision-making in natural intelligence. DSL framework allows better understanding and modeling how individuals in real-world scenarios exhibit a level of adaptability that current AI systems have yet to achieve.

## 1. Introduction

Behavioral research traditionally explores how individuals address the *credit assignment problem* (CAP) - the challenge of attributing “values” to actions based on their effectiveness in achieving rewards (Doya, 1999; Daw et al., 2005; Niv, 2007; Otto et al., 2013; Dolan and Dayan, 2013; Dezfouli and Balleine, 2013; Cushman and Morris, 2015). Typically, these studies assume a stationary agent structure, where an agent adheres to a consistent learning rule and employs a fixed internal representation of its environment. However, this model does not reflect the complexities of real-world behavior, where an individual’s internal environment representation and learning rule can evolve, resulting in more adaptive behavior.

We introduce a Dynamic Structure Learning (DSL) framework designed to capture how agents transition between different internal model structures. In dynamic structure models (Muzy and Zeigler, 2014a; Uhrmacher, 2001; Barros, 1997), changes of structure consist of the addition, deletion or alteration of model components. We extend here this approach to learning systems. Specifically, we define an Agent Structure (AS) as a combination of an internal environment representation (decision graph) and a learning rule. The latter can be described by a Reinforcement Learning (RL) algorithm, which achieves a particular credit assignment (cf. Figure 1a). By constructing all possible AS combinations from a set of reinforcement learning rules and environment representations, we can infer the most likely sequence of ASs for an individual, based on its behavioural observations.

**Figure 1.**
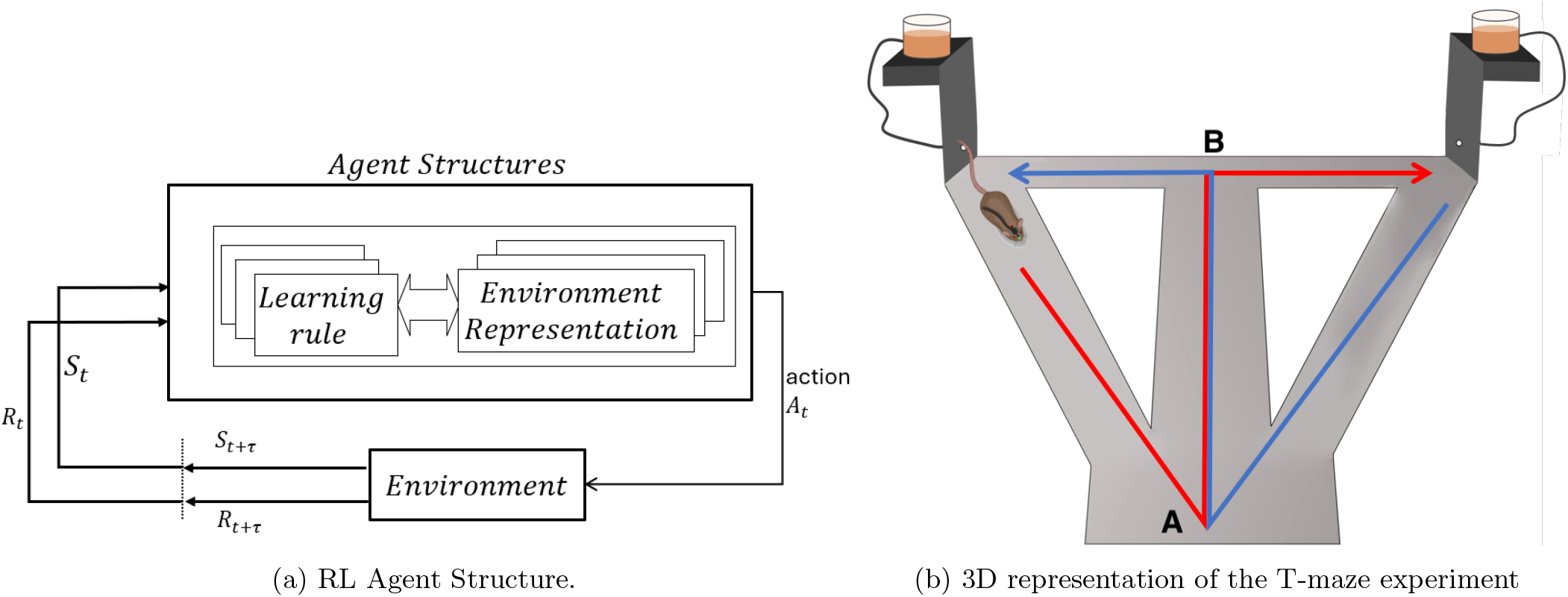
Agent Structure and 3D representation of the T-maze experiment: (a) Semi-Markov Decision Problem (SMDP) formulation of a Reinforcement Learning (RL) problem where agent structure is defined as a combination of learning rule and an internal environment representation, with action of the agent having a random duration *τ* (b) 3D representation of the T-maze experiment: A, B, LF and RF are four choice points. LF and RF represent the Left and Right Feeders respectively. Good path to LF, Good.LF, is shown in red, while good path to RF, good.RF, is shown in blue.

We apply the DSL framework to a T-maze task involving rats, where they must learn the two correct paths between two feeder boxes in the maze (*cf*. , Figure 1b). The rats need to identify the optimal paths leading to rewards in the left feeder (LF) and right feeder (RF). The optimal path from LF differs from that of RF and therefore the optimal behavior in the maze if governed by a hidden rule related to the starting feeder box. Initially, the rats’ Internal Maze Representation (IMR) is suboptimal (*IMR*_*subOpt*_), as they are unaware of the hidden rule and construct an incorrect task representation. Over time, they refine their IMR to match the true optimal decision graph (*IMR*_*opt*_), recognizing the environmental factors (feeder boxes) that influence the reward path, thus reducing errors and increasing reward acquisition.

We also examine two distinct learning rules, achieving distinct credit assignments: Cognitive Activity-based Credit Assignment (CoACA) (James et al., 2023), a heuristic decision-making method based on Activity-based Credit Assignment (Muzy, 2019; James et al., 2023)(ACA), and Q-learning (Watkins and Dayan, 1992), representing a more economically rational approach. CoACA is a “suboptimal learning rule” that reinforces actions based on both rewards and action duration, with longer actions being more memorable (James et al., 2023). This can lead to seemingly irrational choices, as rats may persist with more memorable past actions that offer partial rewards, regardless of their actual economic value. In contrast, Q-learning represents an “optimal learning rule,” aimed at identifying the sequence of actions that maximizes rewards in the maze.

The combination of two learning rules and two environment representations results in four potential agent structures (ASs):

1. The “suboptimal AS”: the combination of the suboptimal learning rule (*LR*_*subOpt*_ or CoACA) and the suboptimal internal maze representation (without feeder boxes) (*IMR*_*subOpt*_),
2. The “LR suboptimal AS”: the combination of the suboptimal learning rule (*LR*_*subOpt*_ or CoACA) and the optimal internal maze representation (with feeder boxes) (*IMR*_*opt*_),
3. The “IMR suboptimal AS”: the combination of the optimal learning rule (*LR*_*opt*_ or Q-learning) and the suboptimal internal maze representation (without feeder boxes) (*IMR*_*subOpt*_), and
4. The “optimal AS”: the combination of the optimal learning rule (*LR*_*opt*_ or Q-learning) and the optimal internal maze representation (with feeder boxes) (*IMR*_*opt*_).

We define rats’ strategies as their ASs, which are given by the combination of learning rules and the internal environment representation they employ. Assuming rats utilize a specific AS to produce behavioral trajectories within a session, our objective is to determine the most probable sequence of these ASs across all sessions. Previously, Inverse Reinforcement Learning (IRL) (Ziebart et al., 2008; Babes et al., 2011; Michini and How, 2012) methods have been used to infer agent’s internal models from observations. IRL methods aim to capture individual behavior by inferring reward functions from observations, with Ashwood et al. (2022) also allowing time-varying reward functions. In Kwon et al. (2020), both reward functions and individual’s beliefs about the world are inferred in a Partially Observable Markov Decision Process (POMDP) setting. Ashwood et al. (2020) present a method for inferring individuals’ learning rules from behavioral data in a perceptual decision-making task, without the need to learn the reward function.

While these inference methods have similarities to our problem, they are typically investigated in settings where state spaces are the same across tasks, differing only in reward functions or transition dynamics. Our approach differs from existing methods to infer agents’ internal models in three key aspects. First, we do not assume that the agent always acts optimally; instead, it can use heuristic decision-making rules, as captured by CoACA. Second, we do not require the agent to have a perfect understanding of the environment; it can operate with a potentially flawed internal representation (*IMR*_*subOpt*_). We also enable the agent to switch between different internal representations, allowing us to model scenarios where, for example, rats might initially use a *suboptimal AS* or *IMR suboptimal AS* based on a simpler yet suboptimal *IMR*_*subOpt*_, and then transition to a more complex but optimal AS based on *IMR*_*opt*_ after learning (see Figure 2).

**Figure 2.**
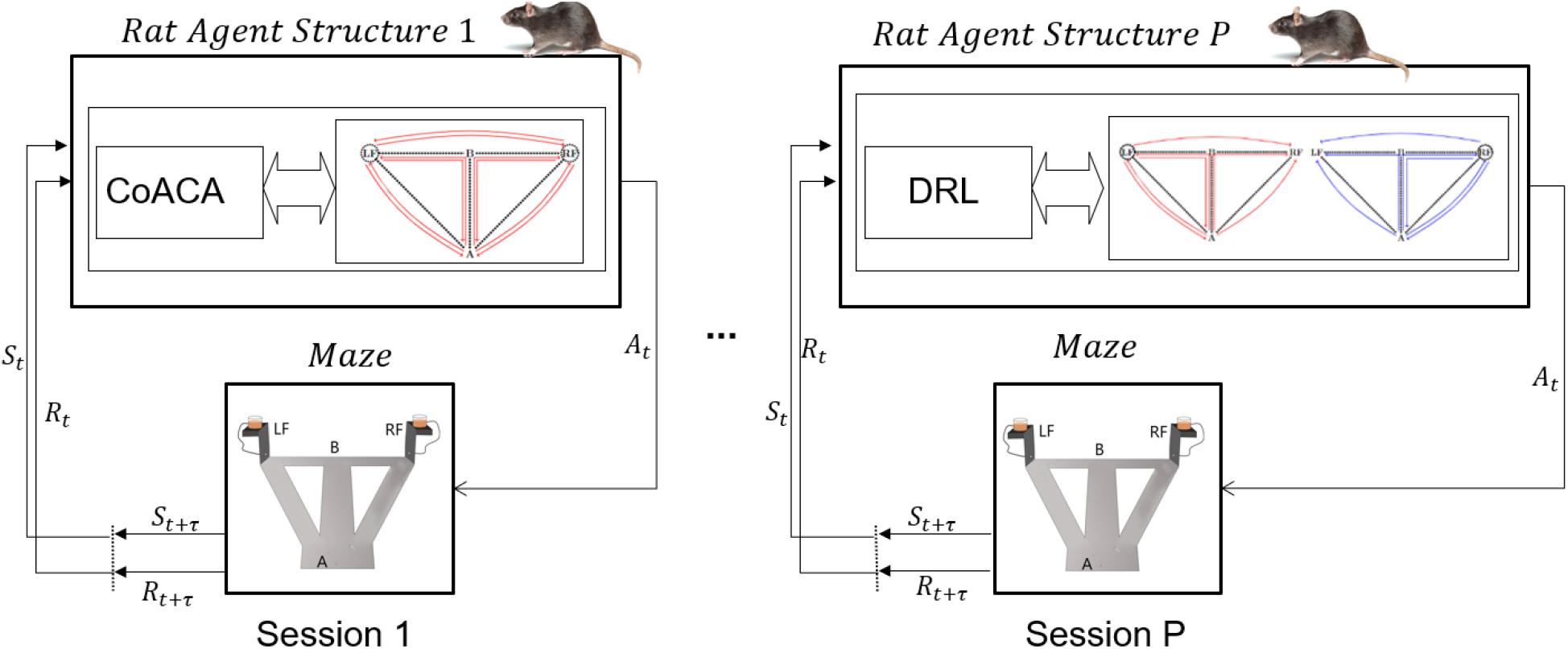
An example of ASs inferred by DSL framework over multiple sessions of rat experiment.

Applying DSL to the rats’ dataset shows that: (i) rats that show slower learning progress appear to rely on the *suboptimal AS* during the early stages of the experiment before switching to the *optimal AS*, whereas rats that learn quickly adopt the *optimal AS* from the beginning of the experiment, (ii) rats’ switches from the *suboptimal AS* to the *optimal AS* indicate a progressive refinement in their perception of the task structure (environment model). The gradual refinement of the IMR requires the rats to “imagine” and construct novel maze representations consistent with their experience, ultimately defining learning as the ability to forge an accurate mental model of the task.

DSL framework offers a more nuanced perspective on learning compared to the traditional dichotomy of exploration and exploitation in reward-based learning (Cohen et al., 2007; Mehlhorn et al., 2015). We demonstrate that randomly exploring the state space to better assess the “value” of actions also includes refining an internal model of the world, transitioning from simpler, flawed model to a more complex, accurate representation of the world. Additionally, the rats’ ability to “imagine” new maze representations underscores a crucial adaptability trait in natural intelligence (Lake et al., 2017). This ability enables them to infer the causal structure of their environment—such as learning the hidden rules that govern the rewards in the maze experiment—and to update their internal models accordingly. In contrast, current AI systems are not capable of inferring causal structures from observations (Lake et al., 2017; Shanahan et al., 2020; Fjelland, 2020; Javed et al., 2020; Bishop, 2021). DSL framework allows better understanding and modeling how individuals in real-world scenarios exhibit a level of adaptability that current AI systems have yet to achieve.

## 2 Methods

### 2.1 Maze experiment

Five male Long-Evans rats were used in the experiment. To motivate the rats to collect food rewards from the maze, they were subjected to a food deprivation program by keeping them at 90% of their body weight during the experiment. Each rat has multiple sessions in the maze, where each session lasts 20 minutes. During the sessions, the rats can freely move around in the maze uninterrupted. The T-maze with return arms (*cf*. Fig. 1b) has two feeder places, Left Feeder (LF) and Right Feeder (RF), where the rats could get a food reward. The maze consists of a central stem (100*cm* long), two choice arms (of 50*cm* each) at one end of the central stem and two lateral arms connecting the other end of the central stem to the choice arms. Prior to the experiment, the rats were trained in the maze for two days, with one 20-minute session per day during which they were free to explore the maze and collect the sugar pellets that were randomly scattered throughout. The experiment began on the third day with two 20-minute sessions.During the sessions, the rats can freely move around in the maze uninterrupted. The maze has four possible choice points, A, B, LF and RF, where the rat must choose between alternative paths. The behavioral task for the rat is to associate the rewards with the two good paths.

### 2.2. Semi-Markov Decision Process

The maze learning task is defined as a Semi-Markov Decision Process (SMDP), which is a generalisation of a Markov Decision Process where actions have a random duration. An SMDP can be defined by a tuple (*S, A, R, T, F* ), where *S* is the set of states, *A* is the set of actions, *R* is the reward function that gives the reward associated with each (*S, A*) in the environment, *T* is the transition function that gives the transition probabilities *Pr*(*s*^*′*^ | (*s, a*)), *F* : *F* (*t* |*s, a*), with *t* ∈ ℝ^+^, gives the probability that next state *s*^*′*^ is reached within time *t* after action *a* is chosen in state *s*.

An *episode* is defined as a minimal segment of the rat’s trajectory where the rat starts from one feeder box, visits the other feeder box and returns to the starting box. Two examples of episode are given below, where 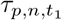 , 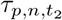 and 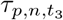 represents the durations of actions taken at times *t*_1_, *t*_2_ and *t*_3_ in episode *n* of session *p*:

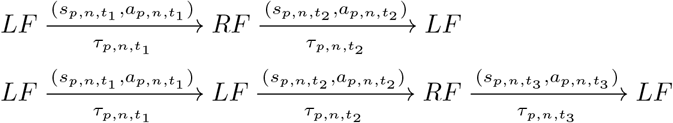

### 2.3. Learning Rules

#### 2.3.1. Cognitive Activity-based Credit Assignment (CoACA)

Cognitive Activity-based Credit Assignment (CoACA) leverages the concept of activity from Activity-based Credit Assignment (Muzy and Zeigler, 2014b; Muzy, 2019). In contrast to traditional RL which views action duration as a cost to minimize, CoACA interprets duration as the effort invested in a choice. This distinction is captured in CoACA’s concept of activity, which acts as a measure of action effort. By prioritizing choices with higher activity (longer duration), CoACA becomes a heuristic decision-making approach – favoring choices that are more memorable due to the effort invested, but not necessarily the most rewarding (James et al., 2023).

Activity is computed as the duration of an action, relative to the duration of an episode:

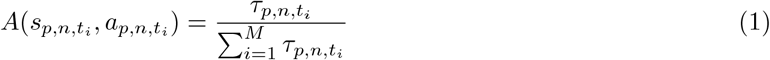

where *t*_*i*_ represents the the time of the *i*^*th*^ action in episode *n* of session *p*, where *i* ∈ [0, *M* ] with M being the total number of actions in the *n*^*t*^*h* episode of *p*^*th*^ session. 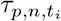 represents the duration of the action taken at time *t*_*i*_ in episode *n* of session *p*.

At the end of a session, the credits of all (*s, a*) pairs in the maze are decayed:

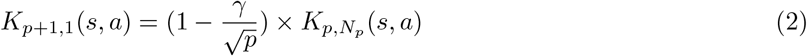

where *γ ∈* [0, 1] is forgetfulness parameter, which decays with time, *i*.*e*., the rats forget less and less with training and *N*_*p*_ represents the final episode of session *p*. Here *t*_*i*_ represents the time at which *i*^*th*^ action of episode *n* in session *p* was taken, *i* ∈ [1, *M* ], *R*_*p*,*n*_ = {0, 1, 2 } is the total reward obtained in episode *n* and *α* is the learning parameter (0, 1]. CoACA implicitly employs memory trace of one episode as it requires the agent to maintain a memory of its choices in the last episode.

The probability of selecting an action *a* in state *s*_*p*,*n*,*t*_ is computed using the softmax rule:

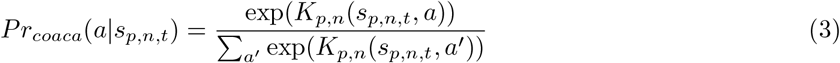

#### 2.3.2. Discounted Reward Reinforcement Learning

A continuous-time version of Q-learning called SMDP Q-learning, which uses temporal difference (TD) errors to iteratively update Q-values, defines the rational behaviour of agents based on an exponential discounting of future rewards (Bradtke and Duff, 1994).

Let 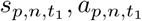 be part of episode *n* of session *p*, leading to new state 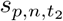 after duration 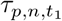 with a reward 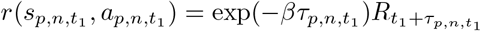 where 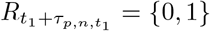 is the reward obtained in the maze after time 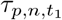 for taking action 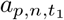 at time *t*_1_, and *β* is the exponential discount factor applied to future rewards. This state transition can be noted as:

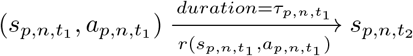

Since CoACA implicitly implements a memory trace of an episode, we implement an eligibility trace in DRL, lasting for the duration of a single episode. At time 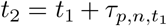 after taking action 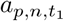 at time *t*_1_, eligibility trace 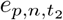 is updated as below:

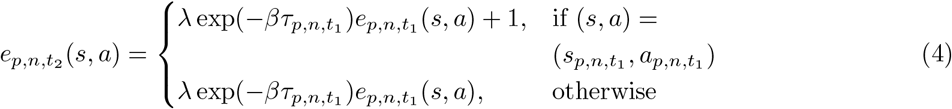

where 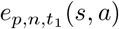 represents the eligibility trace of state-action pair (*s, a*) at time *t*_1_ in episode *n* of session *p*. At the end of an episode, *e*(*s, a*) = 0 *∀*(*s, a*).

Temporal difference prediction error *δ* is given by:

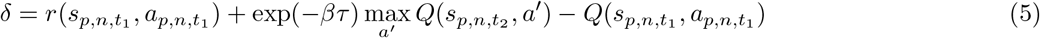

TD update is given by:

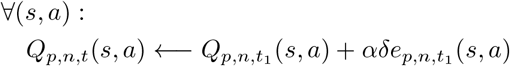

The probability of selection of action *a* in state *s*_*p*,*n*,*t*_ is

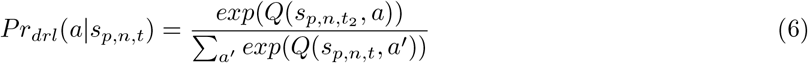

### 2.4. Inferring Rats’ Switching Agent Structures

Our objective is to infer the agent structure (AS) used by the rats in each session based on their experimental trajectories. The AS in session *p* is represented by *x*_*p*_ ∈ {*suboptimal AS, LR suboptimal AS*,*IMR suboptimal AS*,*optimal AS*}. The complete log-likelihood consists of the joint distribution of the unknown ASs *x*_1:*P*_ and the observed trajectories of each session *y*_1:*P*_ , where *P* is the final session, can be expressed as:

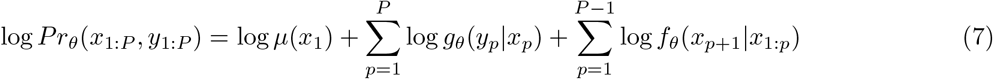

where the initial probabilities *μ*(*x*_1_) are initialized uniformly to 0.25, *g*_*θ*_(*y*_*p*_ | *x*_*p*_) gives the likelihood of observations *y*_*p*_ in *p*^*th*^ session and *f*_*θ*_(*x*_*p*+1_ | *x*_1:*p*_) gives the transition probabilities of ASs given all past ASs and *θ* represents the parameters estimated from rats’ experimental data.

Rats are more likely to employ an AS based on *IMR*_*subOpt*_ early in the experiment, with a shift to *IMR*_*opt*_ based AS once they learn the true structure of the task. To capture this shift in the probability of ASs during the course of the experiment, we utilize a time-varying transition function based on Chinese restaurant process (CRP) (Aldous et al., 2006). This function defines the probability of employing an AS based on its popularity (the number of times it has been chosen previously). The transition function *f*_*θ*_(*x*_*p*_ | *x*_1:*p−*1_) is defined below.

For *k* = 1, 2, 3, 4 representing the four ASs, the occurrences of each of the four ASs in the previous sessions *p −* 1 is given by:

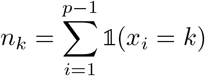

The number of ASs that been chosen at least once until session *p* is given by:

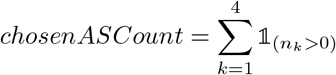

The transition function *f*_*θ*_(*x*_*p*_|*x*_1:*p−*1_) is defined for two scenarios: Case 1, where the AS with *IMR*_*opt*_ has not yet been selected, allowing the rat to explore new ASs, and Case 2, where the AS with *IMR*_*opt*_ has already been chosen, limiting the rat to switching between previously selected ASs without trying any new ones.

**Case 1:** If *optimal AS* or *LR suboptimal AS* has not been selected until session *p*, the probability of selecting AS in session p is given by:

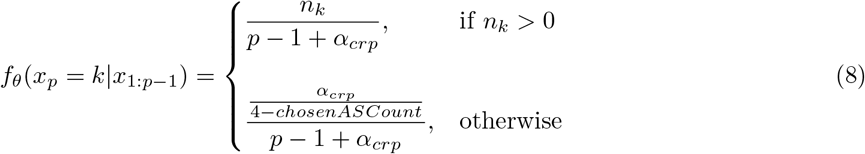

where *n*_*k*_ is the number of times AS *k* has been selected during sessions 1 : *p −* 1, *α*_*crp*_ is the concentration parameter of CRP.

**Case 2:** If either *optimal AS* or *LR suboptimal AS* is selected once:

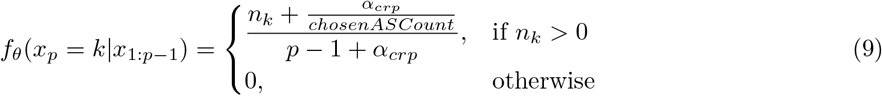

In our study, observations *y*_*p*_ are the trajectories of the rat in a particular session *p* and *g*(*y*_*p*_|*x*_*p*_) gives the probability of trajectory *y*_*p*_ in session *p*:

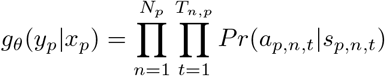

where *N*_*p*_ represents the total number of episodes in session *p* and depending of the value of *x*_*p*_, *Pr*(*a*_*p*,*n*,*t*_ | *s*_*p*,*n*,*t*_) can be given either by Equation (3) or Equation (6).

To infer ASs of rats from their behavioral observations, we employ the Dynamic Structure Learning (DSL) method in Algorithm 1 that computes the smoothing distribution of ASs given by *Pr*(*x*_*p*_ | *y*_1:*P*_ ) and takes the Maximum A Posteriori estimate to determine the AS in each session. Since standard particle filters do not give good estimates of the smoothing distribution, we use Conditional Particle Filter With Ancestor Sampling (CPF-AS) that generates samples from the joint smoothing distribution *Pr*(*x*_1:*P*_ | *y*_1:*P*_ ) (Lindsten et al., 2014).

#### Algorithm 1

Dynamic Structure Learning

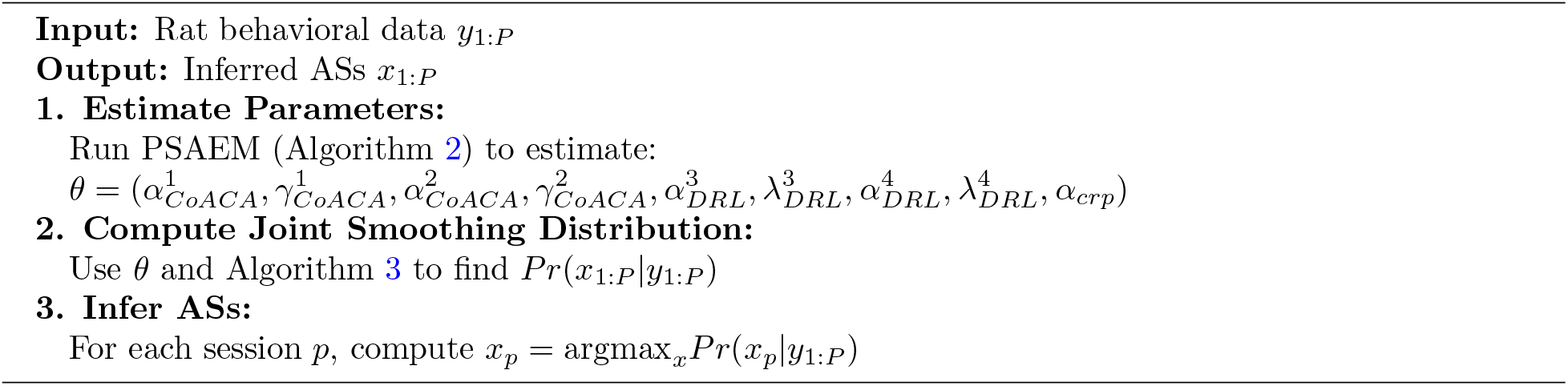

The parameters *θ* of the four different ASs learned from the rat experiment data are:

1. *suboptimal AS* : 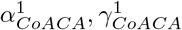
2. *LR suboptimal AS* : 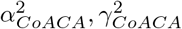
3. *IMR suboptimal AS* : 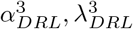
4. *optimal AS* : 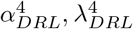
5. CRP concentration parameter: 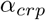

Discount rate 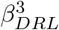 in *IMR suboptimal AS* and 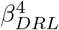 in *optimal AS* are set to 10^*−*4^ so that CoACA and DRL models have two paameters each.

In Algorithm 2, model parameters *θ* are estimated by computing the maximum likelihood estimate by applying Stochastic Approximation Expectation-Maximization (SAEM) to CPF-AS (Lindsten et al., 2013; Lindholm and Lindsten, 2018), which computes the maximum likelihood estimate. Line 8 of Algorithm 2 represents the E-step of SAEM, where 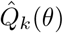 is estimated using Equation 7. In the M-step (Algorithm 2, line 9), new parameters *θ*_*k*_, maximizing the 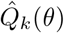 , are determined using the Self-adaptive Differential Evolution optimiser from the Pagmo cpp package (Biscani and Izzo, 2020).

#### Algorithm 2

Particle Stochastic Approximation Expectation Maximization (PSAEM)

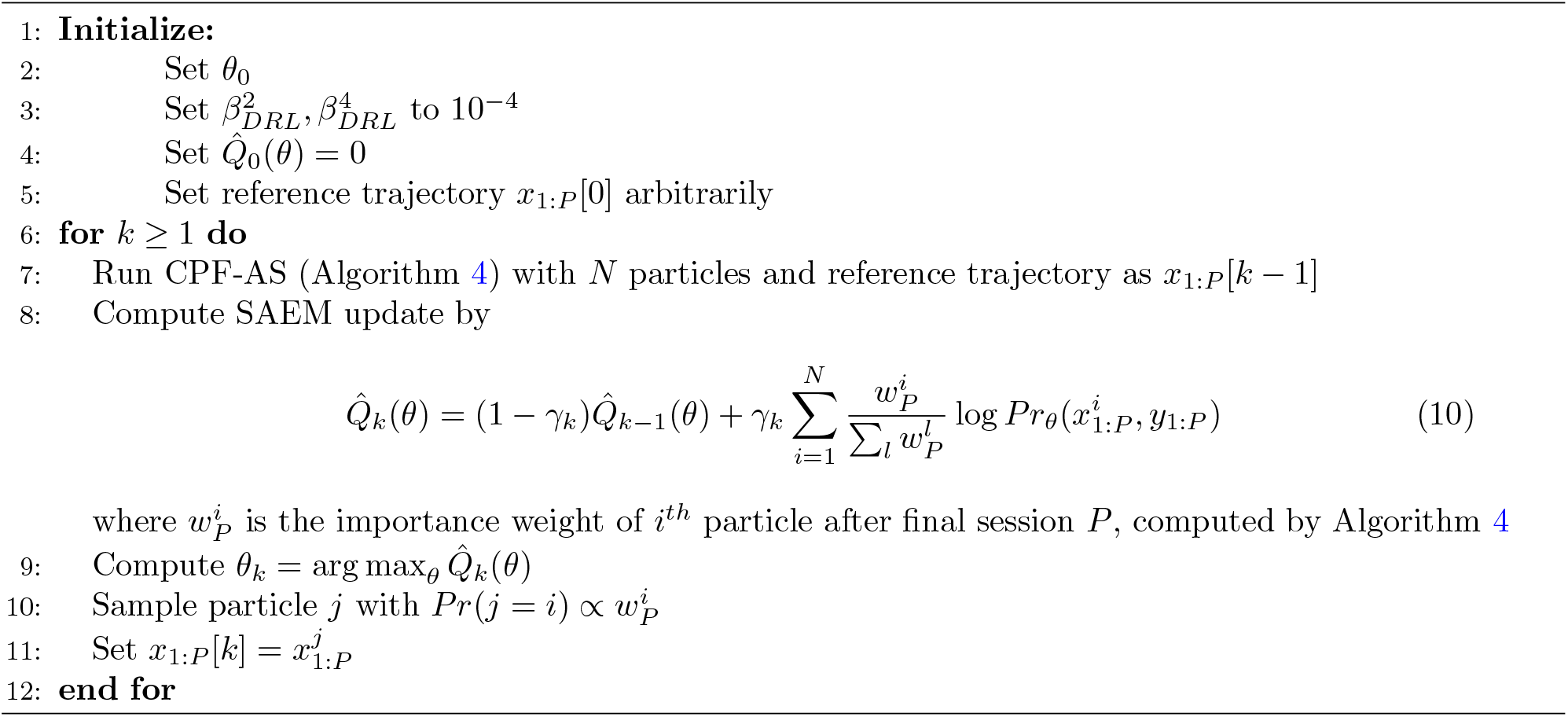

The joint smoothing distribution *Pr*_*θ*_(*x*_1:*P*_ | *y*_1:*P*_ ) is computed using Algorithm 3 and model parameters determined applying Algorithm 2 on experimental data. The most likely sequence of ASs in each session is determined as the Maximum A Posteriori (MAP) estimate of the smoothing distribution *Pr*_*θ*_(*x*_*p*_ | *y*_1:*P*_ ), where *p* represents the current session and *P* is the final session.

To estimate the smoothing distribution, we use the Conditional Particle Filter with Ancestor Sampling (CPF-AS) (Lindsten et al., 2014, 2013). This algorithm takes a reference trajectory *x*_1:*P*_ and outputs a new trajectory 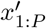 which is a sample from the joint smoothing distribution *Pr*_*θ*_(*x*_1:*P*_ |*y*_1:*P*_ ).

#### Algorithm 3

Smoothing Algorithm

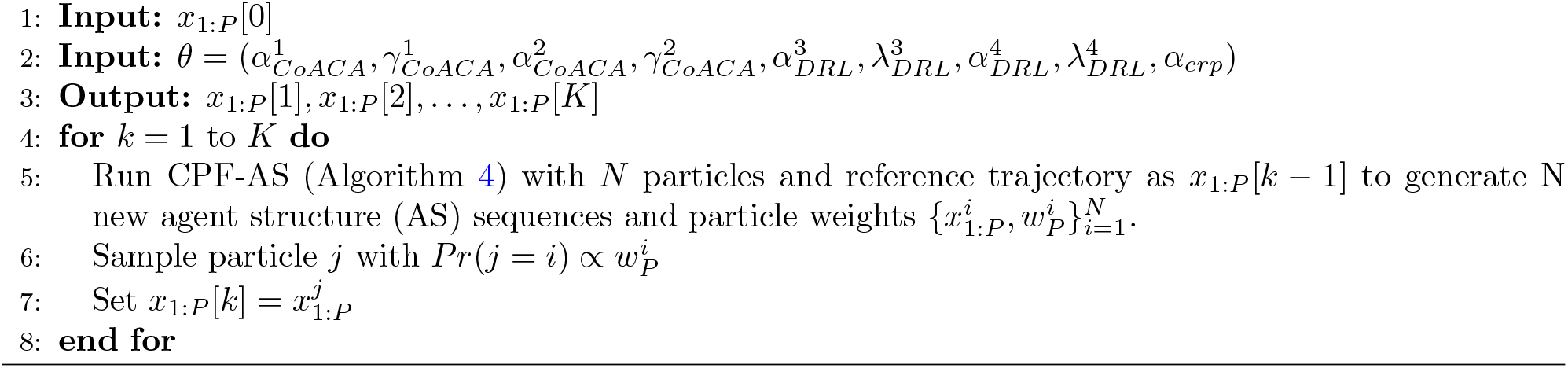

A standard particle filter, based on importance sampling and resampling, approximates the filter distribution *Pr*_*θ*_(*x*_*p*_ | *y*_1:*p*_) by iteratively performing the following steps to update the filtering distribution as new observations arrive:

- Resample: Simulate new ancestors 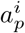 for each particle *i*, according to the importance weights from the previous iteration (line 5 in Algorithm 4).
- Propagate: Particles are propogated to next timestep by sampling from the proposal distribution *r*(*x*_*p*_ | *x*_1:*p−*1_, *y*_*p*_) that incorporates both past ASs *x*_1:*p−*1_ and current observations *y*_*p*_ to generate ASs for session *p*.
- Weight: Compute new importance weights 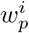 using Equation 13.

While particle filter provides accurate filtering estimates *Pr*_*θ*_(*x*_*p*_|*y*_1:*pS*_), it does not generate good estimates of the joint smoothing distribution *Pr*_*θ*_(*x*_1:*P*_ | *y*_1:*P*_ ) due to the issue of path degeneracy as repeated resampling leads to all particles sharing common ancestors for sessions *p << P* (Chopin et al., 2020). Here we use CPF-AS (see Algorithm 4) to estimate the smoothing distribution, where CPF-AS acts as a Markov Chain Monte Carlo (MCMC) kernel with stationary distribution given by the joint smoothing distribution *Pr*_*θ*_(*x*_1:*P*_ |*y*_1:*P*_ )(Lindsten et al., 2014). We run CPF-AS with *N* = 30 particles. Each particle *i* has an ancestral trajectory 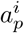 that represents the ASs from sessions 1 : *p −* 1. The ancestral path of each particle represents a potential sequence of ASs, reflecting the behavior of a rat in the maze. Each particle maintains its own unique set of credits or q-values for each of the four different ASs based on its ancestral trajectory 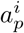 . A locally optimal proposal distribution is used to propagate particles to time *p* given by (Chopin et al., 2020)

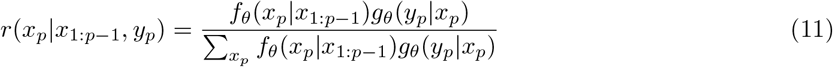

In CPF-AS, the *N*^*th*^ particle ASs 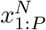 are deterministically set to input reference trajectory. The ancestor of the *N*^*th*^ particle is resampled based on the ancestor weights given by Equation (12). Since the ASs evolve in non-Markovian manner in our models, (Lindsten et al., 2014) provides a a non-Markovian adaptation where the product is truncated to *L* steps, which implies a gradual decay of the non-Markovian influence of the current time step *p* over the next *L* steps. In our analysis we set (*L* = 5).

## 3. Results

### 3.1 Behavioral Analysis

As shown in Fig. 3, the rats trajectories in the initial sessions highlight the differences among slow-learning rats (rat1, rat2 and rat3) and the fast-learning rats (rat4 and rat5). The fast-learning rats learn to get rewards from both LF and RF consistently, whereas the slow learning rats seem to get fewer rewards during the early sessions.

**Figure 3.**
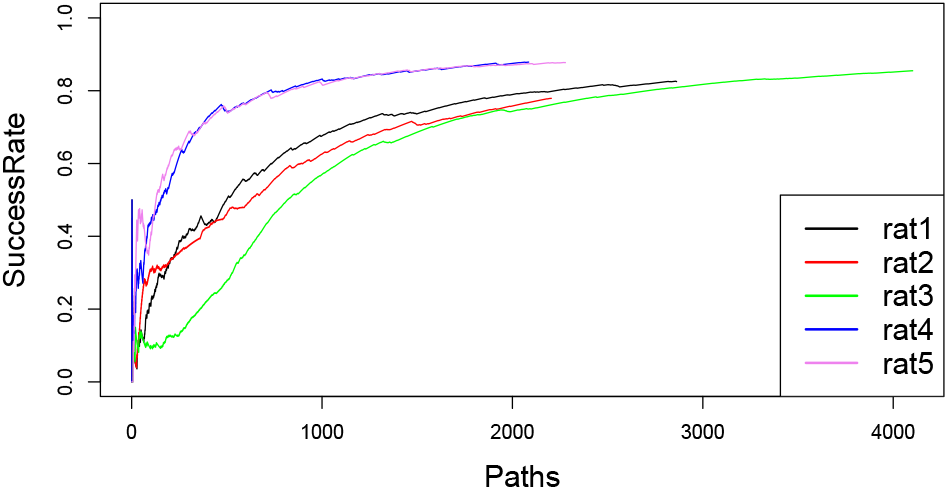
Success rate as proportion of rewarded paths: Success rate computed as a proportion of rewarded paths: The rats can be categorized as slower learning (rat1, rat2, rat3) or faster learning (rat4, rat5) based on the proportion of rewarded paths.

### 3.2. Cognitive Insights Into Rats’ Suboptimal Behaviors

During the learning stage, the rats make significantly more loop path errors compared to other errors. Table 1 shows the number of errors of each type (loop, backward loop, reverse and v) (*cf*. Fig. 4) made by the rats during the learning stage (first 400 paths). *Chi-square test* shows that the looping errors occur above chance levels and cannot be explained as simply random chance events during the learning phase of the rats.

**Table 1:** Error path comparison.

|  | V | Inverse | Loop | Inverted Loop | Are all wrong paths equally likely? |
| --- | --- | --- | --- | --- | --- |
| rat1 | 2 | 1 | 43 | 1 | No ($pval < 2.2 \cdot 10^{-16}$) |
| rat2 | 2 | 0 | 19 | 2 | No ($pval = 6 \cdot 10^{-9}$) |
| rat3 | 5 | 4 | 72 | 4 | No ($pval < 2.2 \cdot 10^{-16}$) |
| rat4 | 8 | 3 | 13 | 5 | No ($pval = 4.9 \cdot 10^{-2}$) |
| rat5 | 6 | 2 | 17 | 4 | No ($pval = 3.3 \cdot 10^{-4}$) |

**Figure 4.**
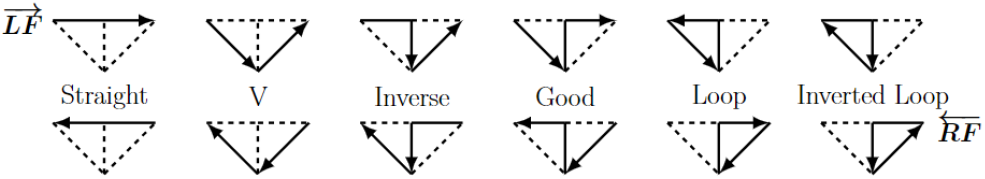
Valid paths in the maze. Turning back in reward boxes are not considered as valid paths as the rats rarely moved backward during the experiment. Top column shows paths starting in Left Feeder (LF) and bottom column shows paths starting in Right Feeder (RF).

#### Algorithm 4

Conditional Particle Filter with Ancestor Sampling (CPF-AS)

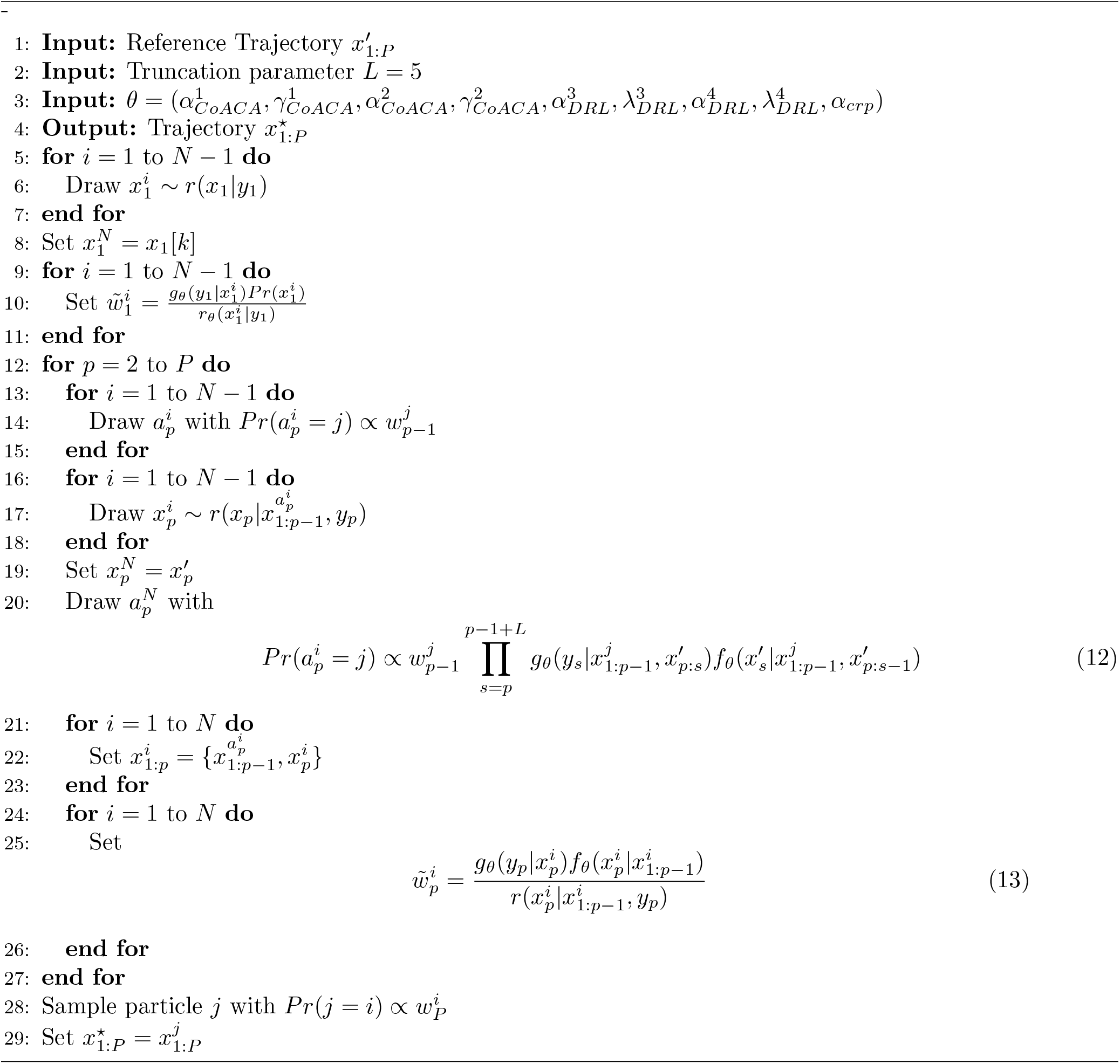

A possible explanation for the high number of loop errors is that the rats might be misinterpreting the reward association. The final segment of their successful path from the Left Feeder (LF) (red dotted line in Figure 5) could be mistakenly linked to the reward itself (located at the Right Feeder, RF). Since both the successful “Good.LF” path and the looping “Loop.RF” path share the segment A *→* B *→* RF, the rats might attempt to replicate this sequence even when starting from RF, hoping to receive another reward (depicted by the blue dotted line in Figure 5).

**Figure 5.**
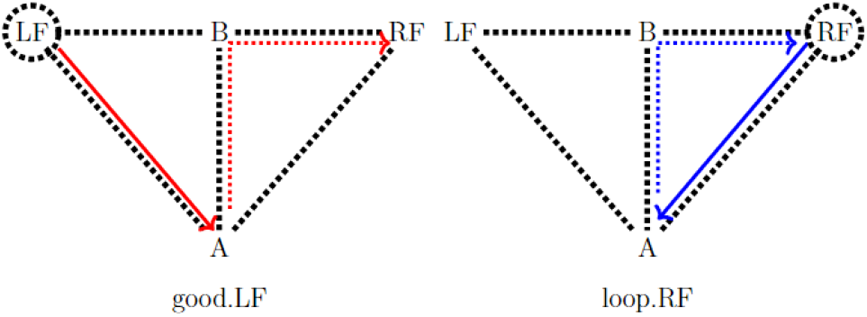
Loop error: rats mistakenly associate the trajectory *A → B → RF* with reward. In suboptimal representation, *A → B → RF* while starting in LF (in good.LF) is same as *A → B → RF* while coming from RF (in loop.RF). Dotted circle indicates the starting feeder box.

Alternate explanations for the high number of loop errors are possible but they are not in agreement with experiment data:

- The loop path could arise because the rats forget which feeder they come from and mistakenly decide to return to the same feeder. If this were the case, then this behavior should consistently persist throughout the experiment.
- It is possible that the rats receive a reward and simply want to revisit the same feeder, anticipating more rewards. However, if their sole motivation were to return to the last feeder, a similar preference for both loop and “inverted loop” (returning directly to LF) would be expected. The data suggest a specific preference for the loop path, indicating a different underlying cause.

Based on the explanation that rats make more loop errors due to mistakenly associating the final segment of the “Good” path with the reward (as shown in Figure 5), we can hypothesize that these loop errors arise because rats are unaware that “starting feeder box” defines the next reward path. Essentially, they ignore the role of this hidden factor and instead repeat the final segment of the previous “Good” path, leading to the last reward.

Building upon this insight, we will proceed to define both a suboptimal and an optimal decision graph in the subsequent section to further understand and characterize the rats’ ASs.

### 3.3. Behavioral Models

#### 3.3.1 Internal Maze Representations (IMR): Suboptimal vs Optimal

Fig. 6a represents *IMR*_*subOpt*_, a suboptimal version of the maze decision graph, not accounting for the *starting feeder box*. Here *A → B → RF* coming from *LF* shares the same representation with *A → B → RF* coming from *RF* , thus leading rats to make loop errors (*cf*. Fig. 5) while searching for rewards.

**Figure 6.**
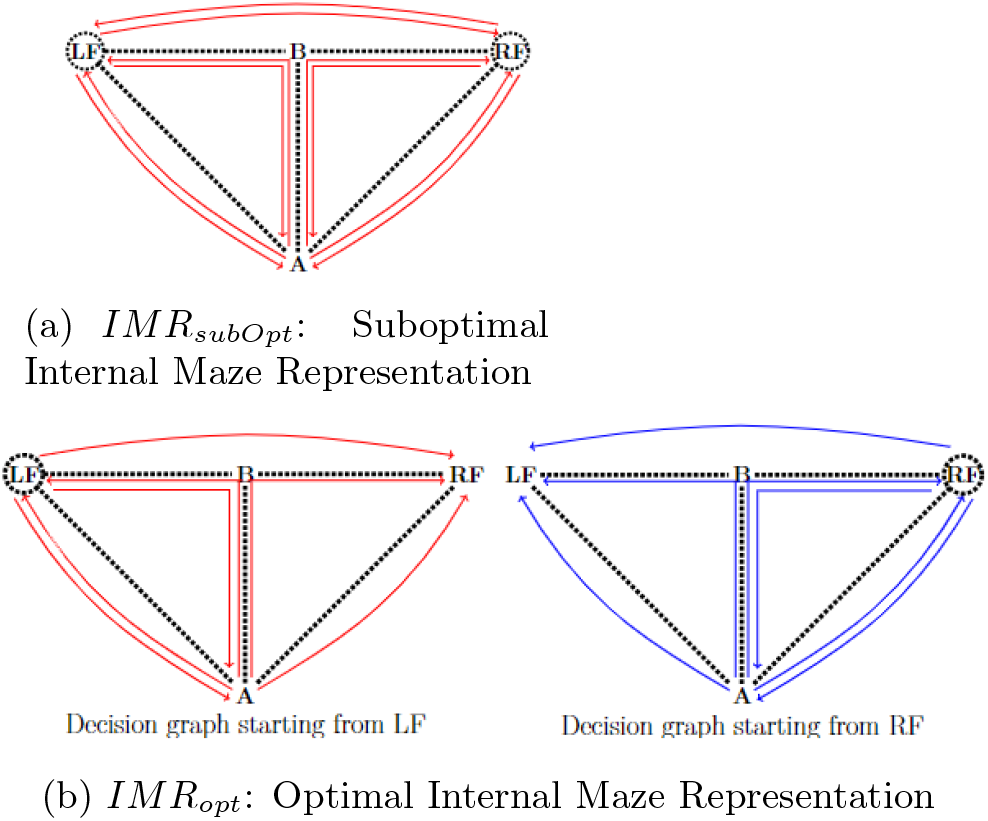
Suboptimal and optimal maze representations: a) *IMR*_*subOpt*_: Suboptimal Internal Maze Representation not accounting for starting feeder box. This implies that trajectories such as *A → B → RF* have the same representation whether they start from LF or RF. b) *IMR*_*opt*_: Optimal Internal Maze Representation with two different decision graphs based on the starting feeder (LF or RF). Here *A →B →RF* coming from LF is different from *A → B → RF* coming from RF. Dotted circles indicate the starting feeder box of trajectories.

The optimal maze representation in the maze task, *IMR*_*opt*_, represented by Fig. 6b, has a larger state space with a separate decision graph for trajectories starting from *LF* and trajectories starting from *RF* . Unlike *IMR*_*subOpt*_, *IMR*_*opt*_, has two separate representations for trajectories such as *A → B → RF* coming from *LF* and *A → B → RF* coming from *RF* , thus having a larger state space and avoiding loop errors in the maze.

#### 3.3.2. Rats’ Agent Structures

We define rats’ AS as a combination of a internal maze representation (*cf*. Fig. 6) and a reinforcement learning rule to solve the *credit assignment problem*. We use two different learning rules (LR):

1. *LR*_*subOpt*_: Cognitive Activity-based Credit Assignment (CoACA) (*cf*. Section 2.3.1)
2. *LR*_*opt*_: Discounted reward RL (DRL), which is a continuous-time version of Q-learning (cf. Section 2.3.2)

Using the two internal maze representations and the two RL methods, we construct four different ASs that the rats could employ in the maze:

- *suboptimal AS* : *LR*_*subOpt*_ with *IMR*_*subOpt*_
- *LR suboptimal AS* : Hybrid AS of *LR*_*subOpt*_ with *IMR*_*opt*_
- *IMR suboptimal AS* : Hybrid AS of *LR*_*opt*_ with *IMR*_*subOpt*_
- *optimal AS* : *LR*_*opt*_ with *IMR*_*opt*_

Considering all four ASs for each session, the total number of combinations for a rat across P sessions would be 4^*P*^ . However, to better reflect reality, we limit the possibilities to six specific ASs that fall into two categories:

1. **Switching from suboptimal to optimal representation:** The rat might start with *IMR*_*subOpt*_, but can still switch to the optimal one later.
2. **Sticking with the optimal representation:** Once a rat chooses a AS with *IMR*_*opt*_, it stays with that choice throughout the experiment.

By focusing on below six possible AS combinations, we create a more realistic model that captures the decision-making switch process of the rats:

1. *suboptimal AS → LR suboptimal AS*
2. *suboptimal AS → optimal AS*
3. *IMR suboptimal AS → LR suboptimal AS*
4. *IMR suboptimal AS → optimal AS*
5. *LR suboptimal AS*
6. *optimal AS*

### 3.4. Inference On Rat Data

To infer how rats switch between agent structures (ASs), we used the Dynamic Structure Learning (DSL) method (see Algorithm 1). This involved first performing model fitting on the experimental data by combining the Conditional Particle Filter with Ancestor Sampling (CPF-AS) (Lindsten et al., 2014) (see Algorithm 4) with Stochastic Approximation Expectation-Maximization (SAEM), following Algorithm 2 (Lindsten, 2013; Lindholm and Lindsten, 2018). The model parameters estimated through Algorithm 2 are presented in Table 2. The agent structures (ASs) for each session were identified by calculating the Maximum A Posteriori (MAP) estimate of the smoothing distribution *Pr*(*x*_*p*_|*y*_1:*P*_ ) determined using Algorithm 3.

**Table 2:** Parameters estimated using Algorithm 2 on experimental data of rats.

| Rats | acaSubopt | | acaOpt | | drlSubopt | | drlOpt | | $\alpha_{crp}$ |
| --- | --- | --- | --- | --- | --- | --- | --- | --- | --- |
| | $\alpha$ | $\gamma$ | $\alpha$ | $\gamma$ | $\alpha$ | $\lambda$ | $\alpha$ | $\lambda$ | |
| rat1 | 0.07 | 0.37 | 0.93 | 0.85 | 0.14 | 0.12 | 0.03 | 0.90 | 1.88 |
| rat2 | 0.32 | 0.56 | 0.26 | 0.92 | 0.73 | 0.88 | 0.07 | 0.43 | 4.03 |
| rat3 | 0.077 | 0.19 | 0.71 | 0.85 | 0.66 | 0.31 | 0.02 | 0.65 | 4.18 |
| rat4 | 0.26 | 0.46 | 0.94 | 0.80 | 0.14 | 0.81 | 0.05 | 0.72 | 1.63 |
| rat5 | 0.78 | 0.06 | 0.59 | 0.96 | 0.52 | 0.05 | 0.05 | 0.52 | 4.15 |

Inference results in Fig. 7 show that the slow learning rats - rat1, rat2 and rat3, utilize the *suboptimal AS* during the initial few sessions before switching the *optimal AS*. In addition, rat1 seems to switch between *suboptimal AS* and *optimal AS*, before settling on *optimal AS*. In contrast, fast learning rats (rat4 and rat5) seem to learn the optimal maze representation early in the experiment and their behaviour is captured by *optimal AS* throughout the experiment.

**Figure 7.**
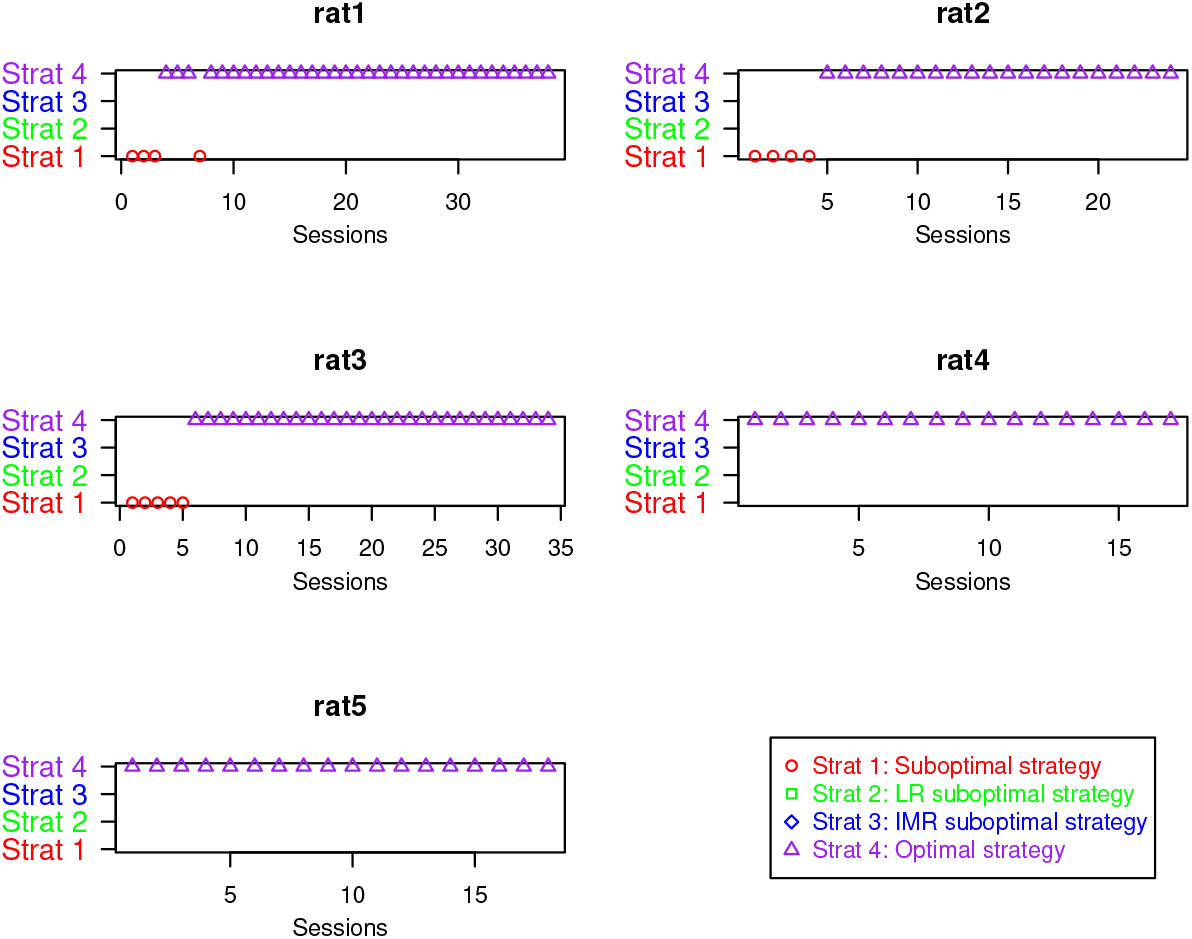
Agent Structures (ASs) of rats inferred using DSL method (see Algorithm 1). ASs result in “strategies” followed by the rats to obtain rewards.

The behaviour of the slow learning rats - rat1, rat2 and rat3 - where they use the *suboptimal AS* in the first sessions leads to a high frequency of loop errors (cf. Table 1) without learning the good path from LF and RF. The fast learning rats, on the other hand, are quicker to use the *optimal AS*, even if they also make loop errors in the beginning (see Table 1). The slow learners showed the cognitive flexibility over time to recognise the need to incorporate “start feeder box” into their internal maze representation and to transition to an optimal behaviour AS. The transition from a suboptimal to an optimal AS over successive sessions highlights two key aspects of the learning process:

1. Ability of rats to “imagine” and adopt a new, more complex internal maze representation that matches their empirical observations.
2. Nature of learning as an ongoing process of refining and improving the internal maze representation.

### 3.5 Simulation Validation

We used simulations to analyze how well DSL recovers the true ASs used to generate the simulated data. We generated simulated trajectories of rats according to the six possible combinations defined in the Section 3.3.2. Parameter recovery on simulated data using Algorithm 2 is plotted as boxplot of the recovery error between the true parameter value and the value recovered from the simulated data is shown in Fig. 8. Overall, the parameter recovery error is small, except in the case of *LR suboptimal AS*. Specifically, the parameter gamma exhibits high variance during recovery. This could be attributed to the fact that gamma represents the decaying forgetting process in Equation 3, which can be replicated by a wider range of parameters.

**Figure 8.**
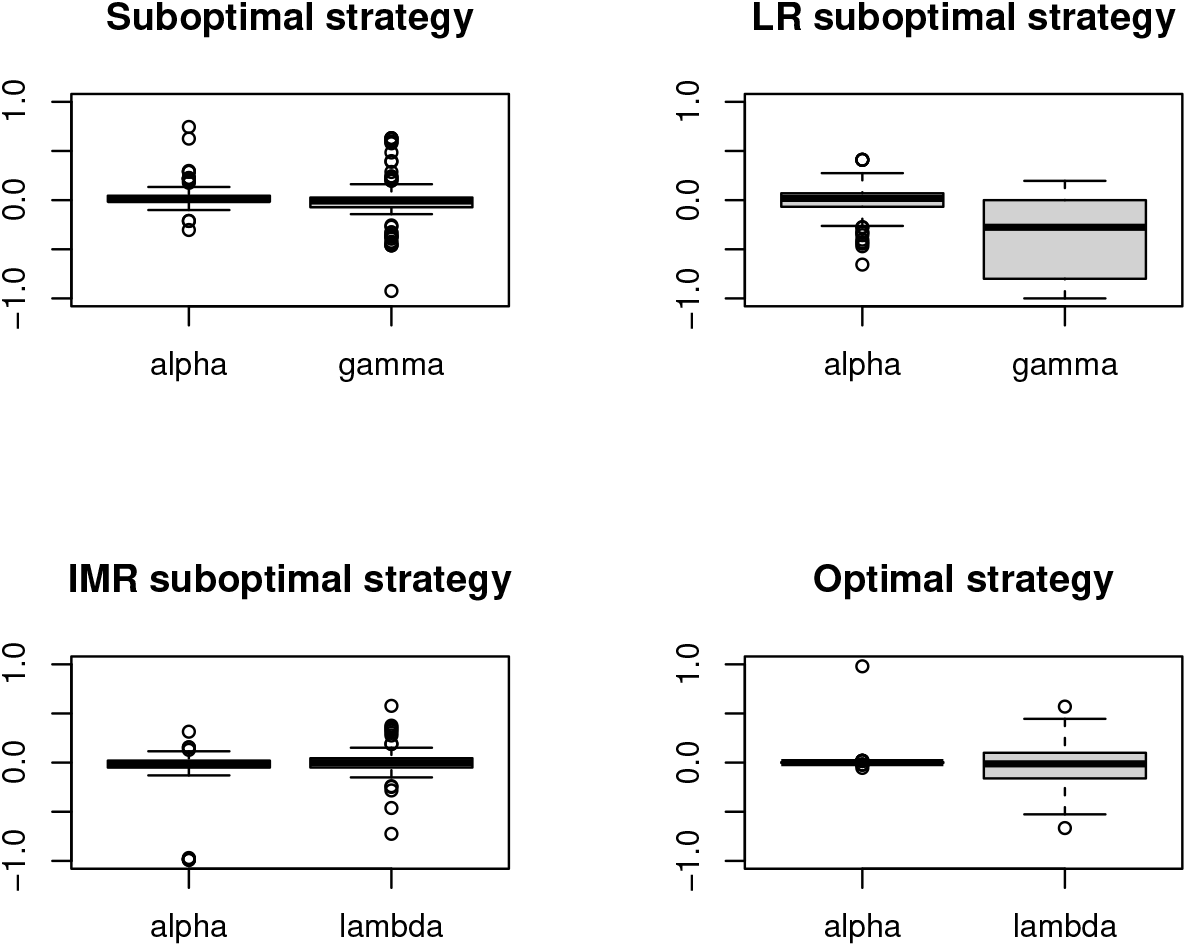
Boxplots showing the errors between the parameter values estimated by the DSL method and the true values on simulated data. The data is based on 300 simulations, with 60 simulations per each of the six possible ASs.

AS recovery is tested by using DSL method (see Algorithm 1) to recover ASs from simulated data. Fig. 9 shows two examples where the true ASs were perfectly recovered. The recovery rate of agent structutres (ASs) across sessions, based on 300 simulations with 60 instances of each of the six possible AS combinations for 5 rats, is shown in Table 3.

**Table 3:** Recovery rate of agent structures (ASs) across sessions for six different AS combinations, determined using the DSL method on simulated data

| True AS | Recovered AS |  |  |  |  |
| --- | --- | --- | --- | --- | --- |
|  | suboptimal AS | LR suboptimal AS | IMR suboptimal AS | optimal AS | None |
| suboptimal AS | 0.90 | 0.09 | 0.01 | 0 | 0 |
| LR suboptimal AS | 0.01 | 0.99 | 0 | 0 | 0 |
| suboptimal AS | 0.96 | 0 | 0.01 | 0.03 | 0 |
| optimal AS | 0 | 0 | 0 | 1 | 0 |
| IMR suboptimal AS | 0 | 0 | 0.96 | 0.04 | 0 |
| LR suboptimal AS | 0 | 0.90 | 0.01 | 0.08 | 0.01 |
| IMR suboptimal AS | 0 | 0 | 0.96 | 0.04 | 0 |
| optimal AS | 0 | 0 | 0 | 1 | 0 |
| LR suboptimal AS | 0 | 1 | 0 | 0 | 0 |
| optimal AS | 0 | 0 | 0 | 1 | 0 |

**Figure 9.**
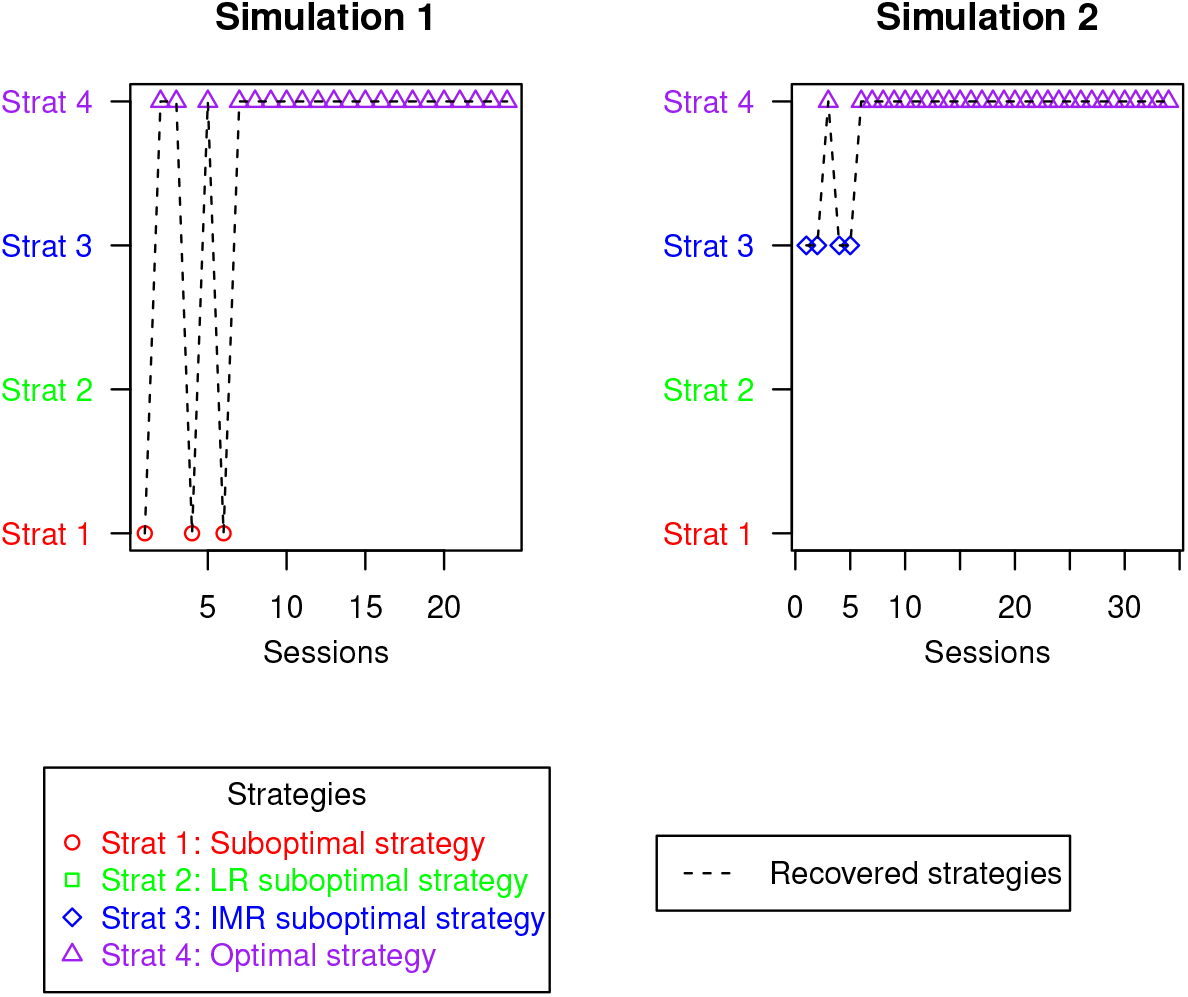
Recovery examples with successful recovery using DSL method on simulated data: Simulation 1 (using rat1 parameters) where AS switches from *suboptimal AS → optimal AS* ; Simulation 2 (using rat3 parameters) where AS switches from *LR suboptimal AS → optimal AS*

## 4. Conclusion

We developed a Dynamic Structure Learning (DSL) framework to reveal the internal model of agents. The latter relies on their evolving cognitive processes behind their behavior. DSL models an agent’s internal dynamics as the interaction between its learning rule and its internal representation of the environment. These core components are drawn from a set of potential candidates, enabling a flexible and adaptive representation of the agent’s decision-making process. The DSL framework aims to reconstruct the most likely sequence of agent structures (ASs) based on observations of the agent, utilizing various learning rules and environment representations.

We applied the DSL framework to the observation from a maze experiment where rats were free to move around without restriction and received rewards in either the Left Feeder (LF) or the Right Feeder (RF) if they took the designated correct paths from LF to RF and *vice versa*. The task required the rats to infer the role of the hidden factor *starting feeder box*, which determined the next reward path in the maze. Thus, the rats could form different possible internal representation of the task, such as one in *starting feeder box* does not influence the observations and another where it influences.

Successfully earning rewards in the task required building a true internal model of the task (incorporating the hidden factors of the task) as well as solving the *credit assignment problem*. Solving both the *internal environment model building* and *credit assignment* problems simultaneously could be a computationally intensive task, with some rats using a suboptimal model of the maze *LR*_*subOpt*_ for a few sessions before eventually switching to the optimal maze model *LR*_*opt*_.

To understand how the rats solved the *internal environment model building* and *credit-assignment* problems, we defined four possible ASs as combinations of two internal maze representations: suboptimal maze model *IMR*_*subOpt*_ and optimal maze model *IMR*_*opt*_; heuristic learning rule *LR*_*subOpt*_ based on CoACA and optimal learning rule *LR*_*opt*_ based on DRL. The *optimal AS*, consisting of *LR*_*opt*_ and *IMR*_*opt*_, represented the ideal AS for the rats to maximise their rewards in the maze. On the other hand, the *suboptimal AS*, composed of *LR*_*subOpt*_ and *IMR*_*subOpt*_, led the rats to potentially suboptimal behavioural patterns, such as a tendency to make loop errors in the maze. We proposed that the loop errors observed in the experimental trajectories of rats during the initial sessions were primarily due to rats employing the suboptimal internal model *IMR*_*subOpt*_, where the reward trajectory overlaps with the loop path, leading rats to mistakenly take the loop path in search of rewards.

The inference results demonstrated that the slow-learning rats appeared to be utilising a *suboptimal AS* in the initial sessions, whereas the fast-learning rats were able to identify the true structure of the task at an early stage of the experiment and appeared to employ *optimal AS* from the initial session. Furthermore, the change in AS from *suboptimal AS* to *optimal AS* in the slow learning rats shows an iterative refinement of their internal maze representations from *IMR*_*subOpt*_ to *IMR*_*opt*_ by the rats.

The slow-learning rats also use a heuristic credit assignment scheme, CoACA, which tends to repeat choices from previously rewarded episodes, with more memorable choices (based on longer duration) receiving higher credits. The rats may rely on a heuristic method either because it is a computationally inexpensive way of repeating choices from memory that were previously involved in a rewarded episode, providing them with partial rewards at the beginning, or because their internal environment representation does not fully explain their observations in the maze.

While our model captures the switching of agent structures by rats throughout the experiment, we assumed that the rats’ internal models are fixed within each session. This approach provides a broader analysis of rat behavior, but a more detailed examination—capable of identifying AS shifts within individual sessions—could offer a deeper understanding of the learning process in individuals.

Our framework to capture the shifting internal structure of agents during their learning provides deeper insights into the learning process of individuals in real-world scenarios. The shift from a suboptimal maze representation, *IMR*_*SubOpt*_, to an optimal one, *IMR*_*opt*_, highlights how the learning process requires individuals to ‘imagine” new possible models of the world when their observations do not match their internal models of the world. This ability of animals and humans to generate previously unexperienced, novel thoughts from past experiences highlights an important aspect of natural intelligence that allows them to adapt to different environments, an ability that remains beyond the scope of AI models. A better understanding of the computational process that drives the “imaginative” process in humans and animals (Buzsáki and Tingley, 2018; Comrie et al., 2022; Kurth-Nelson et al., 2023) could serve to illuminate the capabilities of natural intelligence in flexibily adapting to different environment, paving the way for building more robust AI (Lake et al., 2017; Botvinick et al., 2017; Siemens et al., 2022).

## Acknowledgements

This work was supported by the French government, through the UCA^Jedi^ and 3IA Côte d’Azur Investissements d’Avenir managed by the National Research Agency (ANR-15-IDEX-01 and ANR-19-P3IA-0002), by the interdisciplinary Institute for Modeling in Neuroscience and Cognition (NeuroMod) of the Université Côte d’Azur and by the National Research Agency (ANR-20-CE23-0004) with the DeepSee project. It is part of the Computabrain project.

NEF computing platform from Inria Sophia Antipolis Méditerranée Research Center (CRISAM) has been used for running or parallel simulations. NEF is part of the OPAL distributed computation mesocentre. The authors are grateful to the OPAL infrastructure from Université Côte d’Azur for providing resources and support.

